# Sex-Specific Regulation of Behavioral Responses to Single Prolonged Stress: Role of PACAP

**DOI:** 10.1101/2024.12.11.627918

**Authors:** Marissa A. Smail, Evelin M. Cotella, Susan E. Martelle, James B. Chambers, Ria K. Parikh, Christine E. Moore, Ben A. Packard, Nawshaba Nawreen, Rachel D. Moloney, James P. Herman

## Abstract

Post-Traumatic Stress Disorder (PTSD) is a debilitating condition in which a traumatic experience triggers symptoms related to re-experiencing, avoidance, arousal, and mood dysregulation. PTSD negatively impacts 6% of people during their lifetime, with women being disproportionally affected and exhibiting different, more severe symptoms than men. Despite this widespread impact, the molecular mechanisms underlying PTSD and its sex differences remain poorly understood. Pituitary Adenylate Cyclase-Activating Polypeptide (PACAP) is a neuropeptide which participates in fine-tuning circuitry throughout the brain and has been associated with PTSD in humans, especially in women. Here, we use Single Prolonged Stress (SPS), an animal model of PTSD, to explore the roles of PACAP and sex in PTSD-like behaviors. Specifically, a PACAP agonist or antagonist was infused into the infralimbic (IL) prefrontal cortex, a region key to regulating fear- and anxiety-related behaviors, prior to SPS in male and female rats. One week later, rats were tested in open field/novel object, elevated plus maze, and social interaction. Utilizing a behavioral indexing method, we were able to uncover SPS effects in PTSD-related behavioral domains that were differentially impacted by PACAP manipulations in males and females. While both sexes exhibited increased threat avoidance and decreased threat assessment following SPS, females increased sociability while males decreased sociability. Males also appeared to be protected by IL PACAP antagonism while female SPS phenotypes were exacerbated by IL PACAP agonism. Furthermore, RNAscope revealed that PACAP in the prefrontal cortex responds differently to SPS in males and females. Together, these findings suggest complex relationships between SPS, sex, and IL PACAP which may have important implications for treating PTSD in men and women.

**HIGHLIGHTS:**

- SPS induces different PTSD-like phenotypes in male and female rats
- SPS increases threat avoidance and decreases threat appraisal in both sexes
- Sociability is decreased in males but increased in females following SPS
- IL PACAP manipulation exerts diverging SPS behavioral effects in males and females
- Prefrontal PACAP signaling plays a sex-specific role in SPS molecular mechanisms

## 1. INTRODUCTION

Post-Traumatic Stress Disorder (PTSD) is a detrimental psychiatric condition that impacts an estimated 6% of people in their lifetime^1,2^. PTSD symptoms can emerge following a traumatic event and include intrusive re-experiencing, avoidance, altered arousal, and negative effects on mood and cognition^2,3^. While PTSD has received widespread basic and clinical research attention, its underlying mechanisms remain poorly understood, and the field is still wanting for therapies that can effectively ameliorate PTSD symptoms.

One recently identified candidate for such a therapy is Pituitary Adenylate Cyclase-Activating Polypeptide (PACAP), an excitatory neuropeptide implicated in promoting hormonal and behavioral responses to stress^4,5^. PACAP receptor (PAC1) polymorphisms are associated with risk for development of PTSD^6–10^, and circulating PACAP levels are positively associated with severity of PTSD symptoms^8^. PACAP and its receptors are present throughout many of the brain’s key stress-responsive regions, including the prefrontal cortex (PFC), basolateral amygdala (BLA), central amygdala (CEA), bed nucleus of the stria terminalis (BST), paraventricular nucleus, and hippocampus^4,11,12^. PACAP is generally though to fine-tune this stress circuitry to heighten an animal’s response to environmental challenges^4,5,13^. For example, deletion of the PACAP gene in mice reduces hypothalamic pituitary adrenal (HPA) axis stress responses following various stressors^14,15^. Behaviorally, PACAP knockout mice exhibit increased novelty seeking and reduced threat avoidance behaviors in the elevated plus maze, open field, and light-dark box^16^, consistent with reduced reactivity to novel stressor exposures in the absence of PACAP signaling.

One key circuit in which PACAP may act to modulate PTSD responsivity is the PFC to amygdala connection. PTSD pathology is linked to impaired activation of the ventromedial prefrontal cortex (vmPFC)-amygdala circuits promoting inhibition of fear^17,18^. It is thought that PTSD reduces top-down control of the BLA by the vmPFC, reducing BLA-mediated inhibition of the central amygdaloid nucleus (CEA), resulting in enhanced activation of CEA neurons that drive fear responses^19^. Studies in rodents reveal that central infusions of PACAP disrupt acquisition of fear conditioning, whereas infusions of a PAC1 antagonist disrupts cue learning in a trace conditioning paradigm^20^. A recent study from our group noted long-term enhancement of passive avoidance learning following injection of PACAP into the rodent vmPFC equivalent (infralimbic cortex (IL)), suggesting that the peptide is able to potentiate fear memories via direct modulation of this cortical circuitry^12^. PAC1 receptors in the IL are primarily located on inhibitory interneurons, suggesting that these behavioral effects may stem from PACAP driving inhibition within the IL, impairing its ability to subsequently inhibit amygdala-driven fear responses^12^. In the BLA itself, infusion of PACAP *in vitro* activates neurons projecting to the CEA, thereby inhibiting consolidation of contextual fear conditioning^21^. Together, these data suggest that PACAP can act within vmPFC-amygdala circuitry to exaggerate conditioned fear. The question which remains is how the actions of PACAP within this circuit directly participate in PTSD pathophysiology.

Many of the human PTSD-PACAP associations discussed above are either driven by or present exclusively in women^6–10^. These studies suggest that there are likely sex differences in the interplay between PTSD and PACAP that could have important consequences for the therapeutic potential of this neuropeptide. Despite these disproportionate effects on women, many animal studies of PTSD either focused on males alone or failed to elicit behavioral phenotypes in females. For example, single prolonged stress (SPS), a widely accepted animal model of PTSD, elicits predictable fear-, arousal-, and anxiety-related phenotypes in males^3,22^. SPS in females is much less studied and has yielded more subtle, less consistent behavioral phenotypes^23–25^. Recent studies in our lab have revealed female SPS effects on reinstatement of conditioned fear, but not on more classically studied fear acquisition and extinction-related endpoints that are impacted in males^17^. Despite these behavioral differences, females do exhibit unique molecular changes following SPS^17^. Sex differences in in PACAP expression are noted rodent studies, with females showing higher PACAP tone in the BST and prelimbic cortex (PL)^4,20^.

Here, we utilized SPS in tandem with intra-IL pharmacological manipulations of PACAP to explore 1) the behavioral consequences of SPS in male and female rats, 2) the impact of PACAP agonism and antagonism within the IL on SPS behavioral phenotypes, and 3) the interaction of sex and IL PACAP on these outcomes. Furthermore, RNAscope was used to examine how SPS impacts PACAP in the PFC, providing clues for how these interactions may be governed on different levels. The present results offer insight into the roles of sex and vmPFC PACAP signaling in mediating an individual’s response to a traumatic experience, deepening our understanding of the molecular mechanisms underlying PTSD and offering important considerations for the therapeutic treatment of this detrimental condition.

## 2. MATERIALS AND METHODS

### 2.1 Subjects

Eight-week-old male and female Sprague Dawley rats were purchased from Envigo. All animals were pair-housed in translucent polycarbonate shoebox cages with corncob bedding and food and water available *ad libitum*. Animals receiving cannulation surgery (below) were single housed post-operatively to limit damage to cannulae or cannula implant site. The colony room was temperature and humidity-controlled and maintained on a 12-h light-dark cycle (07:00 h lights on, 19:00 h lights off). All experiments complied with the National Institutes of Health Guidelines for the Care and Use of Animals and approved by the University of Cincinnati Institutional Animal Care and Use Committee.

### 2.2 Experimental Design

Experiment 1 was designed to assess the interaction of SPS and IL PACAP manipulations. Male and female rats received intra-IL infusions of either vehicle (VEH), a PAC1 receptor agonist PACAP 1-38 (PACAP), or a PAC1 receptor antagonist PACAP 6-38 (ANTAG) 30 min prior to SPS. Behavioral assessments began one week after SPS (Fig 1A). Experiment 2 examined the impact of SPS on PFC PACAP. Behaviorally naïve rats were sacrificed one week after SPS and RNAscope was used to examine PACAP expression in the IL and prelimbic (PL) PFC (Fig 1B). Males and females were run in separate cohorts^26^.

**Figure 1:**
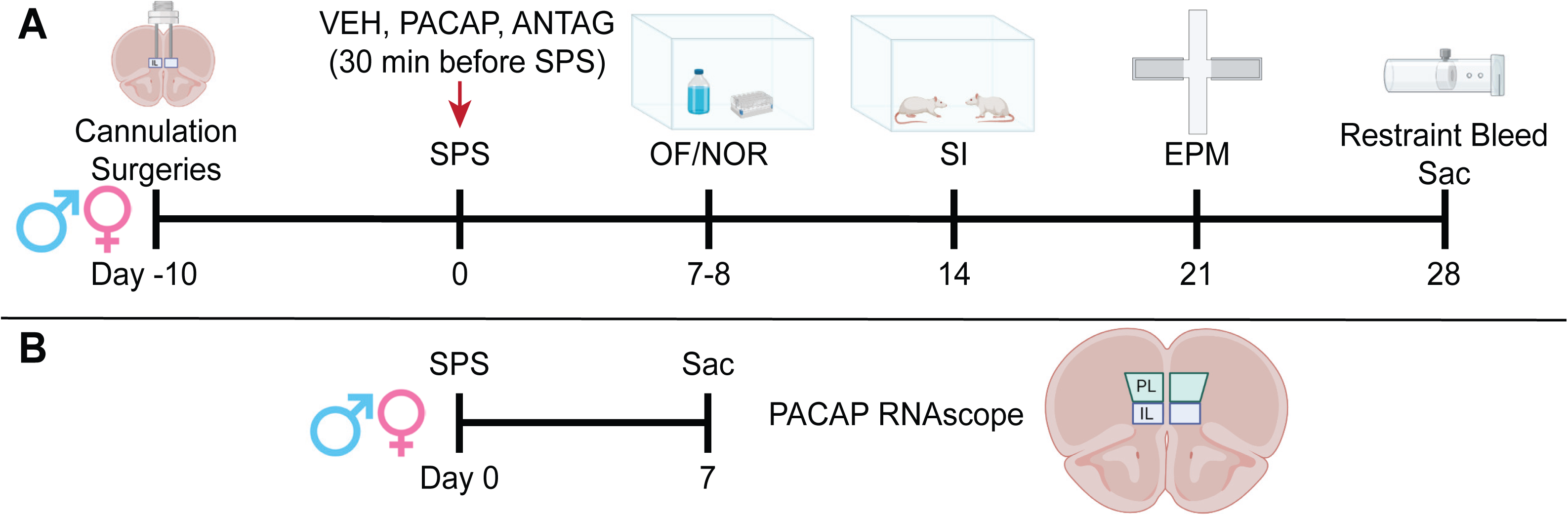
Experimental Design. (A) Experiment 1 Timeline. Male and female rats were implanted with bilateral cannulae targeting the IL. Following recovery, rats were infused with either VEH, PACAP, or ANTAG 30 minutes prior to either SPS or CON. Behavioral testing commenced one week later and consisted of OF/NOR, SI, and EPM, each run a week apart. Rats were then sacrificed 90 minutes after a 30-minute restraint, and brains were collected for verification of cannula placement. (B) Experiment 2 Timeline. Male and female rats were exposed to either SPS or CON, and sacrificed one week later. Brains were collected, frozen, and used for RNAscope analysis of PACAP mRNA expression in the IL and PL. Note that animals in this study were kept behaviorally naïve.

### 2.3 Surgical Procedures

Male and female rats (n = 8-12 per group) were anesthetized with isoflurane and treated with a prophylactic dose (2 mg/kg) of meloxicam (Loxicom®; Norbrook, Overland Park, KS). A stereotaxic instrument (David Kopf Instruments, Tujunga, CA) was used to guide placement of double barreled 26-gauge guide cannulas (Plastics One Inc., Roanoke, VA) into the left and right IL cortices [from bregma: +3.0 mm anterior-posterior, ± 0.6 mm mediolateral, −4.6 dorsoventral] under aseptic conditions. Guide cannulas were fixed with dental cement (Stoelting Company) to 3 anchoring screws on the skull. Each cannula was fitted with a dummy cannula that extended 0.2 mm beyond the tip of the guide. Rats were treated with meloxicam for 2 days following surgery and allowed to recover for 1 week prior to behavioral studies. The dummy cannula was manipulated daily during this period to maintain patency. The locations of the cannula tracks were histologically verified at the end of experiments.

### 2.4 Pharmacological Infusions

PACAP and ANTAG were obtained from BACHEM (Torrence, CA) and dissolved in saline vehicle (VEH) on the day of injection. Thirty minutes prior to SPS, rats received a bilateral intra-IL injection of VEH, PACAP (1µg/500nl), or ANTAG (1µg/500nl) at a rate of 160 nl/min for 3 min via a 33-gauge infusion cannula connected by polyethylene tubing to a 10 μl Hamilton micro- syringe (Hamilton Company). The injectors were left in place for 60 s after injection to allow for diffusion away from the cannula. These doses were chosen because they have been shown previously to be behaviorally active upon intraparenchymal infusion in rats^27,28^. Rats were returned to their home cage until SPS commenced 30 minutes later.

### 2.5 Single Prolonged Stress

SPS was conducted an accordance with prior published procedures^17^. Rats were placed in a plastic restrainer for 2 hours, followed immediately by a 20-minute group swim (n = 6 to 8 animals per tub, 45 cm in diameter, water temperature 25±2°C). Following a 10-minute rest, rats were placed in pairs into a glass desiccator containing ether until unconscious, removed to a clean cage until regaining consciousness, and then returned to the housing room. Animals were left to recover for 1 week before behavioral testing.

### 2.6 Behavioral Testing

#### Open Field and Novel Object Recognition

One week after SPS, rats were subjected to open field (OF) and novel object recognition (NOR) testing. Testing tests consisted of 4 phases: open field (day 1), familiarization 1 (day 1), familiarization 2 (day 2), and novel object testing (day 2). All phases were conducted in a black 75 x 75 x 40 cm plastic box and recorded from above. On the morning of day 1, rats were placed in the center of the empty box and allowed to freely explore for 10 minutes for the open field phase. Approximately 2 hours later, two objects (500 mL bottles filled with blue dye) were placed in the middle of the box, and rats were allowed to freely explore the objects for the 10- minute familiarization 1 phase. The next morning, rats were placed back in with the same 2 objects for the 10-minute familiarization 2 phase. Four hours later, one of the objects was changed to a 12 x 12 x 7 cm tube holder, and rats were allowed to freely explore both objects for the 10-minute novel object phase. Novel and familiar object locations were counterbalanced within each group. Ethovision 13 was used to assess center duration, border duration, and distance moved for the open field phase, as well as time spent exploring (nose within 2 cm) the novel and familiar objects during the novel object phase. A discrimination index was calculated using the formula: (novel duration – familiar duration) / (novel+familiar duration).

### Social Interaction

One week later, rats were subjected to social interaction (SI) testing. Rats were placed in a clean cage (40 x 20 cm) with a novel age- and sex- matched conspecific for 10 minutes. Videos of the interactions were scored by a blinded observer for total time conducting social (e.g., touching, sniffing), non-social (e.g., exploring the cage), aggressive (e.g., pinning), and submissive (e.g., presenting underside) behaviors.

### Elevated Plus Maze

One week later, rats were subjected to elevated plus maze (EPM) testing. The EPM apparatus consists of a 10 cm^2^ center area, two open arms (50 x 10 cm), and two enclosed arms (50 x 10 x 30 cm). Rats were placed one of the open arms, facing the center, and allowed to freely explore the apparatus for 5 minutes. Ethovision 13 was used to assess time spent in the center, open arms, and closed arms, as well as the distance traveled. Additionally, a blinded observer scored the videos for nose exploration (poking the head into the open arms) and nose dips (dipping the head off of the open arms). It is important to note that, for the males, the EPM was run in white light, resulting in little exploration of the open arms. For that reason, the females were then run in dim light, resulting in greater time on the open arms. Differences between males and females on these endpoints thus represent procedural differences rather than differences by sex.

### HPA Axis Assessment

One week later, rats were restrained in plastic restrainers for 30 minutes, and blood was sampled by tail bleed 0, 15, 30, 60, and 120 minutes from the commencement of restraint for corticosterone (CORT) analysis.

### 2.7 Corticosterone Radioimmunoassay

Blood samples were centrifuged (4°C at 6000 rpm for 15 min), and plasma was collected and stored at −20°C until assay preparation. Plasma CORT concentrations were determined using 125I radioimmunoassay kits (MP Biomedicals). All samples were run in duplicate.

### 2.8 Histological Analysis

Following the final tail bleed, rats were administered an overdose of sodium pentobarbital and transcardially perfused with 0.1M phosphate buffer (PBS) followed by 4% paraformaldehyde in 0.1M PBS, pH 7.4. Brains were post-fixed in 4% paraformaldehyde at 4 °C for 24h, then transferred to 30% sucrose in 0.1M PBS at 4 °C until the moment of tissue processing. Brains were sliced into serial 35μm coronal sections using a freezing microtome (−20 °C). Sections (1/6) were collected into wells containing cryoprotectant solution (30% Sucrose, 1% Polyvinyl- pyrolidone (PVP-40), and 30% Ethylene glycol, in 0.1M PBS). Brains were stained with cresyl violet, and the PFC was imaged to assess cannula placement. The correct placement in the IL PFC was determined using the Paxinos & Watson Rat Brain Atlas^29^.

### 2.9 Behavioral Indexing

In order to gain further insights into the present data, we utilized a z-score method to analyze multiple behaviors together and generate indices in behavioral domains known to known to be negatively impacted in PTSD^30^. Z-scores were calculated with the following formula: (x - AVG) / SD, where x = sample value, AVG = average of CON VEH group, and SD = standard deviation of CON VEH group. Z-scores from multiple behavioral endpoints in related domains were then averaged together to generate behavioral indices as follows (Fig 5A). The Threat Avoidance Index included OF Border Duration and EPM closed arm duration. The Threat Appraisal Index included EPM nose exploration duration and EPM head dip duration. The Sociability Index included SI social interaction duration and the negative transform of non-social interaction duration. We additionally correlated these indices together in order to determine if more severe impairments in one domain were indicative of impairments in another.

### 2.10 RNAscope Analysis of Prefrontal PACAP

A new cohort of rats was generated (CON/SPS, n=6/sex/group) and sacrificed by rapid decapitation one week after SPS (Fig 1B). The goal of this study was to determine SPS and sex effects on PFC PACAP expression. Brains were collected, flash frozen in cold isopentane, and stored at -80°C. Brains were warmed to −20°C for 1h and sectioned at 18µm on a Microm cryostat. Sections containing IL and PL PFC were selected for Multiplex RNAscope (Advanced Chemical Diagnostics). The PL was selected as an additional analysis region because of prior studies implicating it in PTSD-related sex differences^20^. Tissue was labeled with a probe for PACAP RNA *adcyap1*. Tissue was fixed in 4% PFA (room temp) for one hour, dehydrated though an ethanol series, and stored in 100% EtOH at –4°C for between 24 hours and a week before staining. Staining was done following the Advanced Chemical Diagnostics multiplex RNAscope protocol for fresh-frozen tissue. Images were obtained bilaterally for the IL and PL (AP +0.5 to +1.5) using a Stellaris confocal microscope and processed using the ACD Hi-plex software. Image settings were kept constant for all animals. PACAP puncta were quantified using Imaris by outlining points in the appropriate channel that matched the diameter of a selected punctum. These data are presented as puncta per cell.

### 2.11 Statistics

It is important to note that we elected to analyze males and female independently in these studies. This decision was made because 1) males and females were assessed in different testing sessions and 2) large inherent male/female differences exist in many of the tested endpoints which could occlude the ability to examine individual endpoints within sex. Therefore, all results were analyzed within each sex and do not directly assess sex differences per se.

Normality and variance were assessed, and two-way (Stress x Infusion) ANOVAs (GraphPad Prism 9) were used to assess behavioral endpoints within sex. Outliers were determined by values that fall outside the mean ± 1.96 times the standard deviation. All post hoc testing utilized Sidak’s multiple comparison tests, which were selected *a priori* to focus on stress effects within infusion groups. Pearson correlations were used where appropriate. CORT curves were analyzed using Two-way repeated measures ANOVAs within sex. RNAscope endpoints were analyzed using unpaired t-tests. Detailed statistical results are provided in Supplementary Table 1.

## 3. RESULTS

### 3.1 Anatomical Verification of Cannula Implantation

Only animals with both cannulae correctly hitting the IL were included in behavioral analyses. Final n’s were as follows: Male CON VEH = 12, Male CON PACAP = 7, Male CON ANTAG = 8, Male SPS VEH = 8, Male SPS PACAP = 7, Male SPS ANTAG = 8, Female CON VEH = 10, Female CON PACAP = 10, Female CON ANTAG = 10, Female SPS VEH = 10, Female SPS PACAP = 9, Female SPS ANTAG = 10.

### 3.2 Impact of SPS and IL PACAP Manipulations on Behavior

SPS decreased OF center duration in males (main effect of stress: F(1,41)=4.197, p=0.047) and females (main effect of stress: F(1,53)=6.107, p=0.017) (Fig 2A,B), suggesting increased anxiety-like behavior. While PACAP manipulations did not significantly impact this endpoint, it appears that ANTAG infusion was able to block this effect in males alone. Border duration was inversely affected, exhibiting increases in both sexes after SPS (male main effect of stress: F(1,41)=4.197, p=0.047; female main effect of stress F(1,53)=6.095, p=0.017); whereas distance moved was not altered in males but exhibited an interaction effect in females (interaction effect F(2,52)=4.382, p=0.017) (Supplementary Fig 1A).

**Figure 2:**
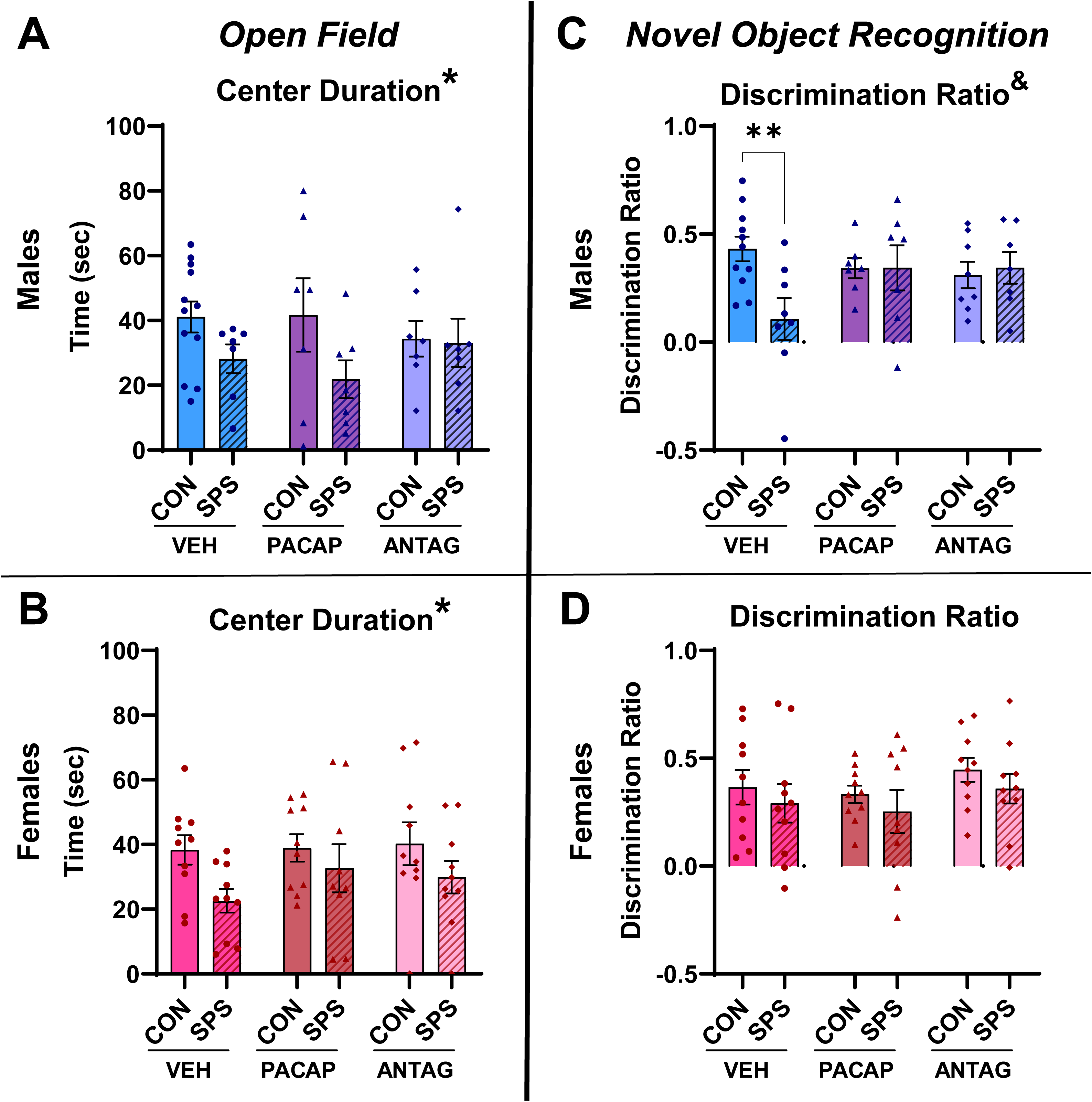
SPS and PACAP Effects on Open Field and Novel Object Recognition Testing. (A) Male OF center duration. (B) Female OF center duration. (C) Male NOR discrimination ratio. (D) Female NOR discrimination ratio. n = 7-12. **\*** = Main Effect of Stress. **&** = Interaction. * = post hoc p<0.05.

NOR exhibited an interesting difference by sex, in that SPS disrupted discrimination in males (interaction effect: F(2,41)=3.733, p=0.032) but not in females (Fig 2C,D). This effect was driven by enhanced exploration of the familiar object in males (interaction effect: F(2,41)=3.917, p=0.028). Females did explore the novel object less following SPS (main effect of stress: F(1,51)=4.195, p=0.046); however, this was not sufficient to impact the overall discrimination ratio (Supplementary Fig 1B). PACAP and ANTAG did not significantly impact discrimination ratio in either sex.

In the SI test, males exhibited decreased social interaction (main effect of stress: F(1,41)=4.081, p=0.049) and increased non-social interaction (main effect of stress: F(1,42)=4.847, p=0.033) following SPS (Fig 3A). These effects were strongest in the VEH animals and largely normalized in both PACAP and ANTAG conditions. Females did not show SPS or PACAP effects on these endpoints (Fig 3B). Females did exhibit stress (main effect of stress: F(1,48)=6.898, p=0.011) and interaction (interaction effect: F(2,48)=4.125, p=0.022) effects on aggressive interactions; however, the low frequency of these events makes it difficult to draw conclusions from these data (Supplementary Fig 1C).

**Figure 3:**
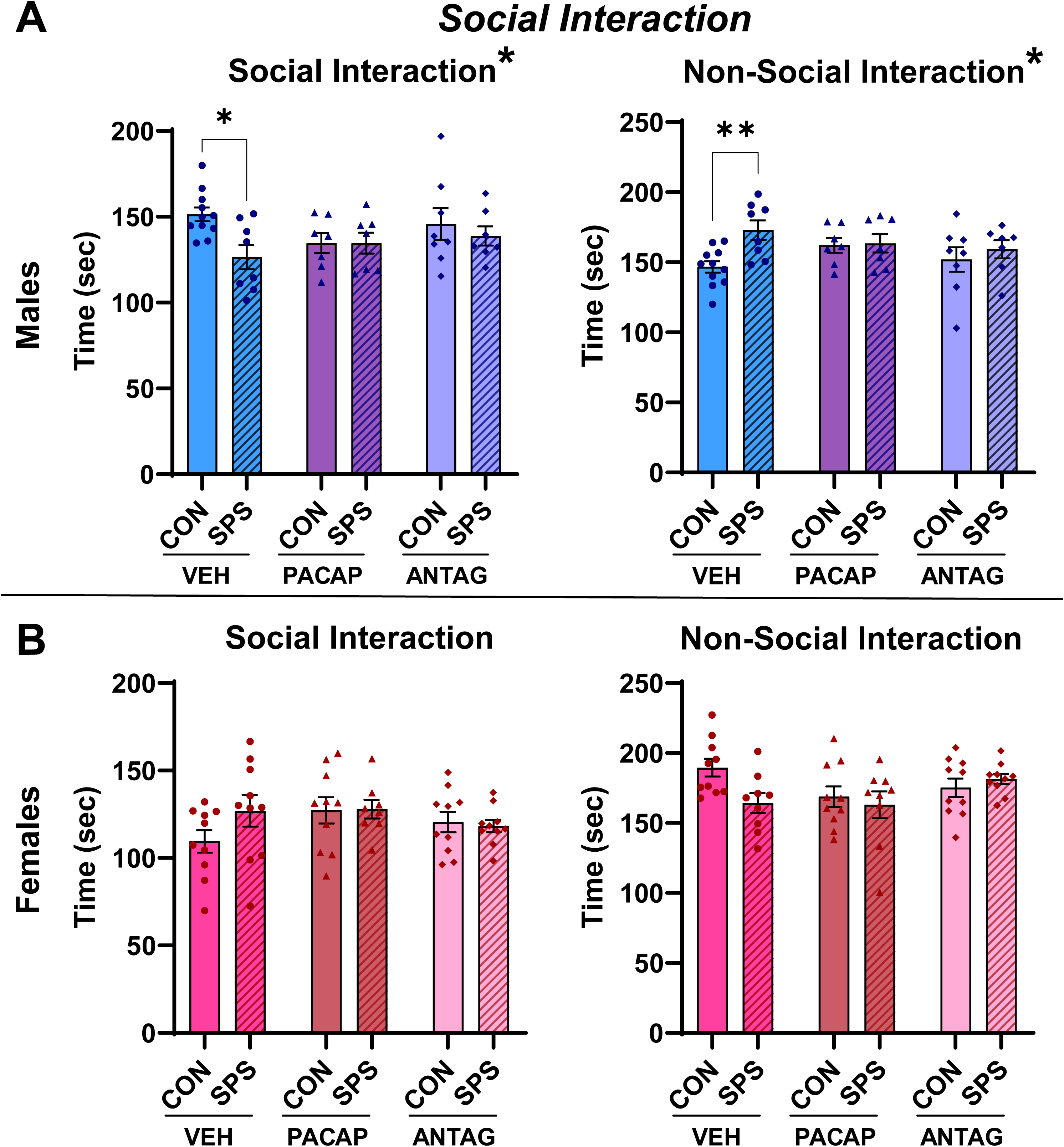
SPS and PACAP Effects on Social Interaction Testing. (A) Male SI social and non-social interaction duration. (B) Female SI social and non-social interaction duration. n = 7-12. **\*** = Main Effect of Stress. * = post hoc p<0.05.

Both sexes exhibited SPS effects on EPM endpoints. In males, closed arm duration was increased by SPS (main effect of stress: F(1,44)=4.634, p=0.037), and center duration was decreased (main effect of stress: F(1,43)=4.580, p=0.038) (Fig 4A). Note that males largely did not go out onto the open arms due to brighter light conditions (Supplementary Fig 1D). This effect was evident in both VEH and PACAP rats, but largely normalized in the ANTAG group.

**Figure 4:**
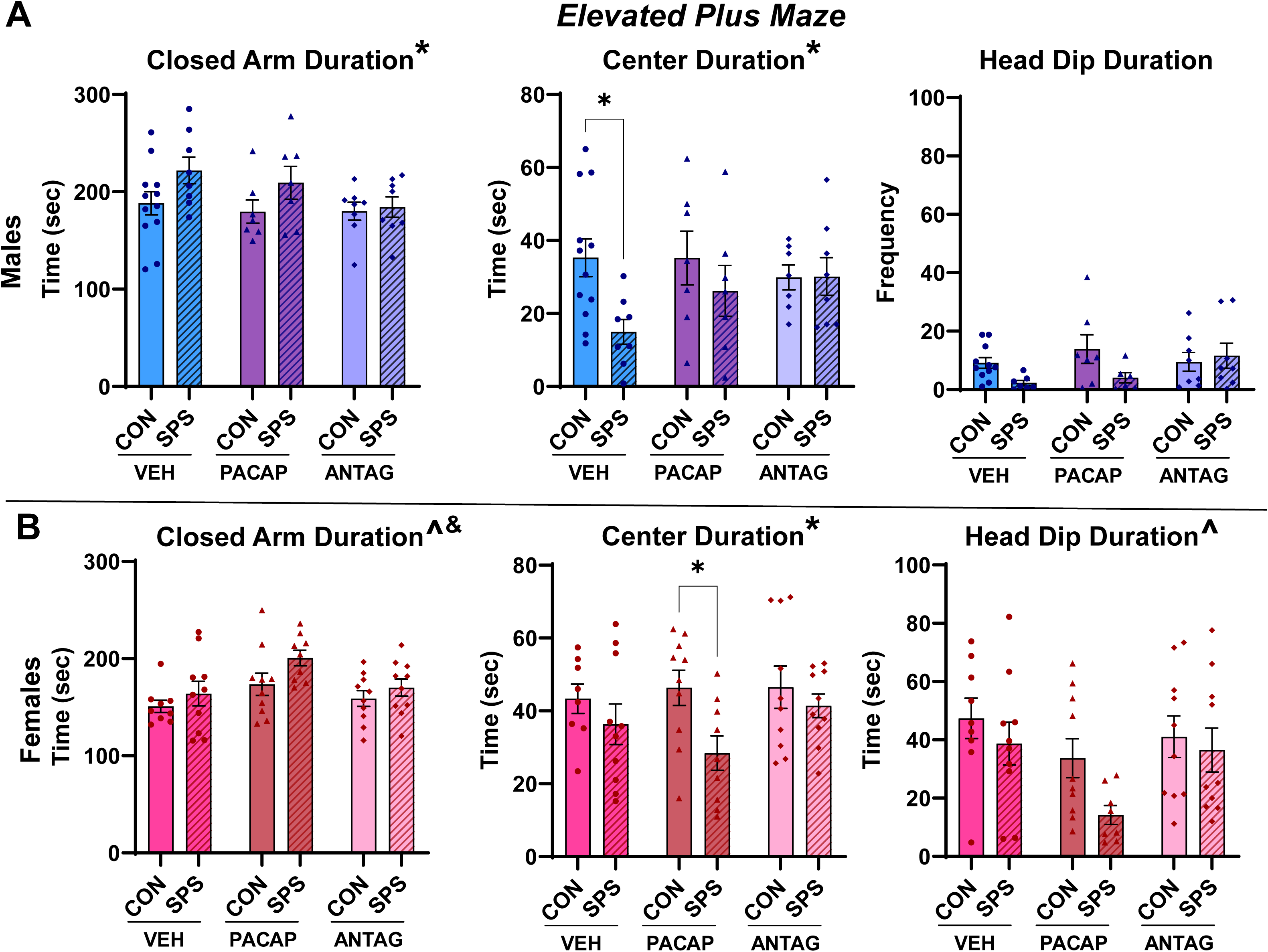
SPS and PACAP Effects on Elevated Plus Maze Testing. (A) Male EPM closed arm duration, center duration, and head dip duration. (B) Female EPM closed arm duration, center duration, and head dip duration. n = 7-12. **\*** = Main Effect of Stress. **^** = Main Effect of Infusion. **&** = Interaction. * = post hoc p<0.05.

Males did not show any significant differences in head dip or nose exploration durations. While females exhibited similar patterns, with SPS increasing closed arm duration (main effect of stress: F(1,52)=4.692, p=0.035; main effect of infusion: F(2,52)=5.069, p=0.010) and decreasing center duration (main effect of stress: F(1,51)=6.338, p=0.015), they appear to have been more affected than males by PACAP infusion (Fig 4B; Supplementary Fig 1D). SPS increased closed arm duration and decreased center and open arm durations the most in females that received PACAP. Similar effects were seen in the other groups, but to a smaller magnitude. Females also exhibited decreased head dip durations (main effect of infusion: F(2,51)=4.155, p=0.021) following SPS, suggesting decreased exploratory behaviors, again driven by the PACAP group.

The CORT response to restraint following SPS and PACAP manipulation was not impacted in either sex (Supplementary Fig 2).

### 3.3 Behavioral Indexing Reveals SPS and PACAP Effects on Complex Behaviors

While the standard behavioral endpoints presented in 3.2 revealed some interesting phenotypes, considering multiple endpoints at once has the potential to offer deeper insight into the interaction of SPS and IL PACAP. The Z-score method was used to generate such combined endpoints, each representing a behavioral domain known to be impacted by PTSD^30^ (Fig 5A).

**Figure 5:**
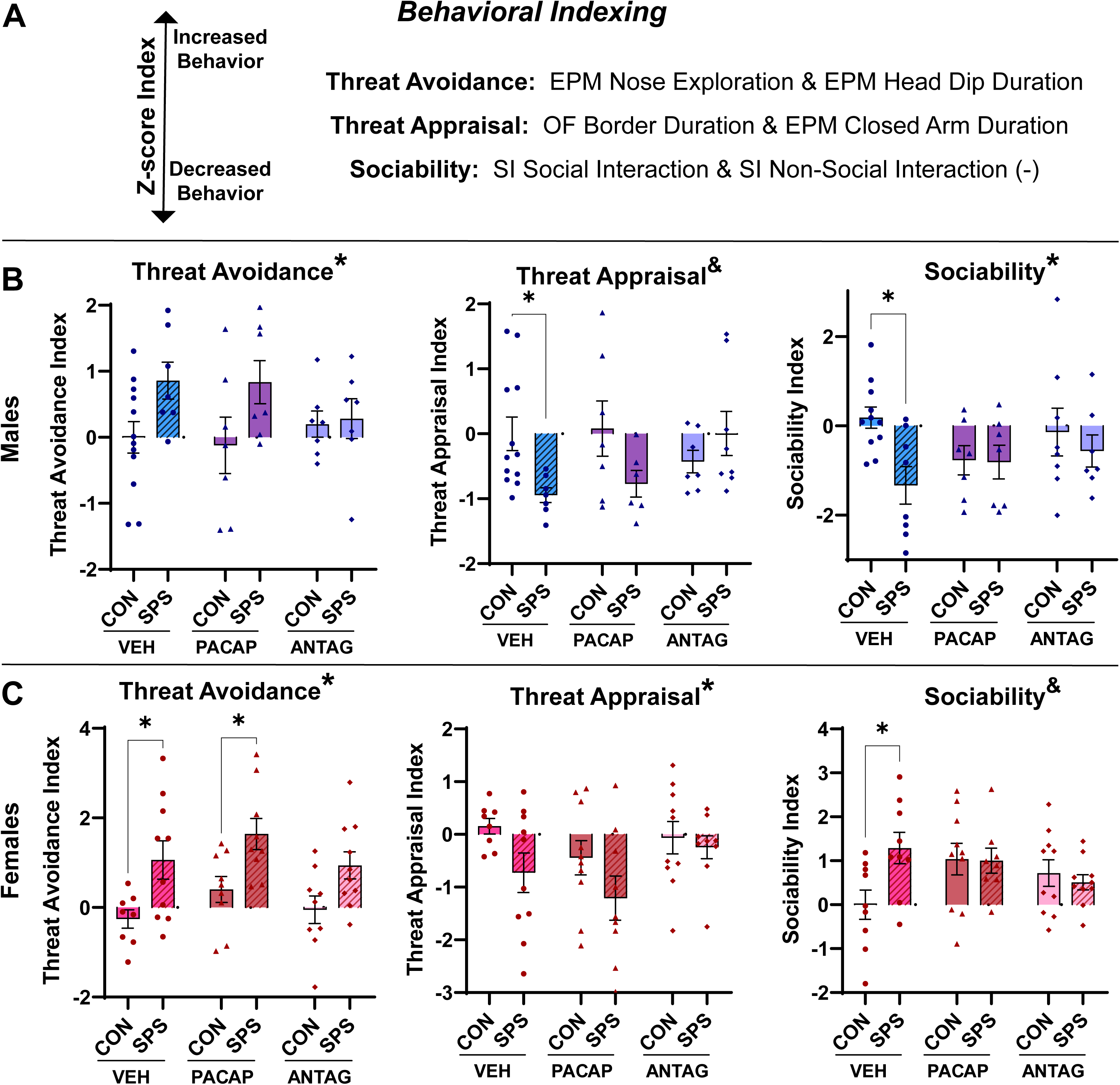
SPS and PACAP Effects on Behavioral Indices. (A) Explanation of behavioral indexing. Z-scores were used to combine behavioral measures into PTSD-relevant indices of threat avoidance, threat appraisal, and sociability. A higher Z-score indicates an increase in that behavioral domain. (B) Male behavioral indices. (C) Female behavioral indices. n = 7-12. **\*** = Main Effect of Stress. **&** = Interaction. * = post hoc p<0.05.

SPS enhanced threat avoidance and decreased threat appraisal in both males (avoidance main effect of stress: F(1,41)=6.509, p=0.015; appraisal interaction effect: F(2,41)=3.580, p=0.037) and females (avoidance main effect of stress: F(1,49)=19.29, p=<0.0001; appraisal main effect of stress: F(1,50)=5.381, p=0.025) (Fig 5B,C). In males, PACAP infusion did not alter these endpoints. However, ANTAG infusion blocked SPS effects on both indices. On the other hand, PACAP moderately enhanced these SPS phenotypes in females, while ANTAG had less of an impact. Thus, while SPS effects on threat-related indices were similar in males and females, the response to PACAP manipulation differed on the basis of sex.

The sociability index also differed by sex. While males exhibited decreased sociability following SPS (main effect of stress: F(1,42)=4.498, p=0.040), females exhibited enhanced sociability (interaction effect: F(2,50)=3.490, p=0.038). Opposing effects were seen in each sex with PACAP and ANTAG, although drug infusion normalized this difference in CON and SPS rats.

Correlating these behavioral indices offers further insight into individual differences in SPS coping strategies (Supplementary Fig 3). In both males and females who experienced SPS, there was a strong negative correlation between threat avoidance and threat appraisal (male R^2^=0.603, p<0.001; female R^2^=0.382, p=0.001), suggesting that as rats became more avoidant, they also become less exploratory of their surroundings, indicating more severe SPS phenotypes. Females also exhibited a negative relationship between threat avoidance and sociability (R^2^=0.161, p=0.042) that was not present in males. This suggests that females who were more social were less avoidant, indicating less severe SPS phenotypes. PACAP manipulations did not significantly shift these relationships.

### 3.4 SPS Differentially Impacts PFC PACAP RNA in Males and Females

A new cohort of rats was generated to assess the impact of SPS on PACAP mRNA expression in the PFC. Brains were collected one week following SPS and RNAscope was used to measure PACAP expression. In the IL, SPS decreased PACAP puncta per cell in male rats (t(66)=3.888, p<0.001) (Fig 6A). Females showed no effect of SPS in this region (Fig 6B). Prior studies have implicated the PL in SPS phenotypes, especially in females, so we also assessed PACAP expression in this region. Here, SPS increased PACAP expression in both sexes (male t(38)=2.396, p=0.022; female t(66)=3.888, p<0.001) (Fig 6C,D). While this effect was stronger in females, it is interesting that the PL may play a role in SPS PACAP interactions in both sexes.

**Figure 6:**
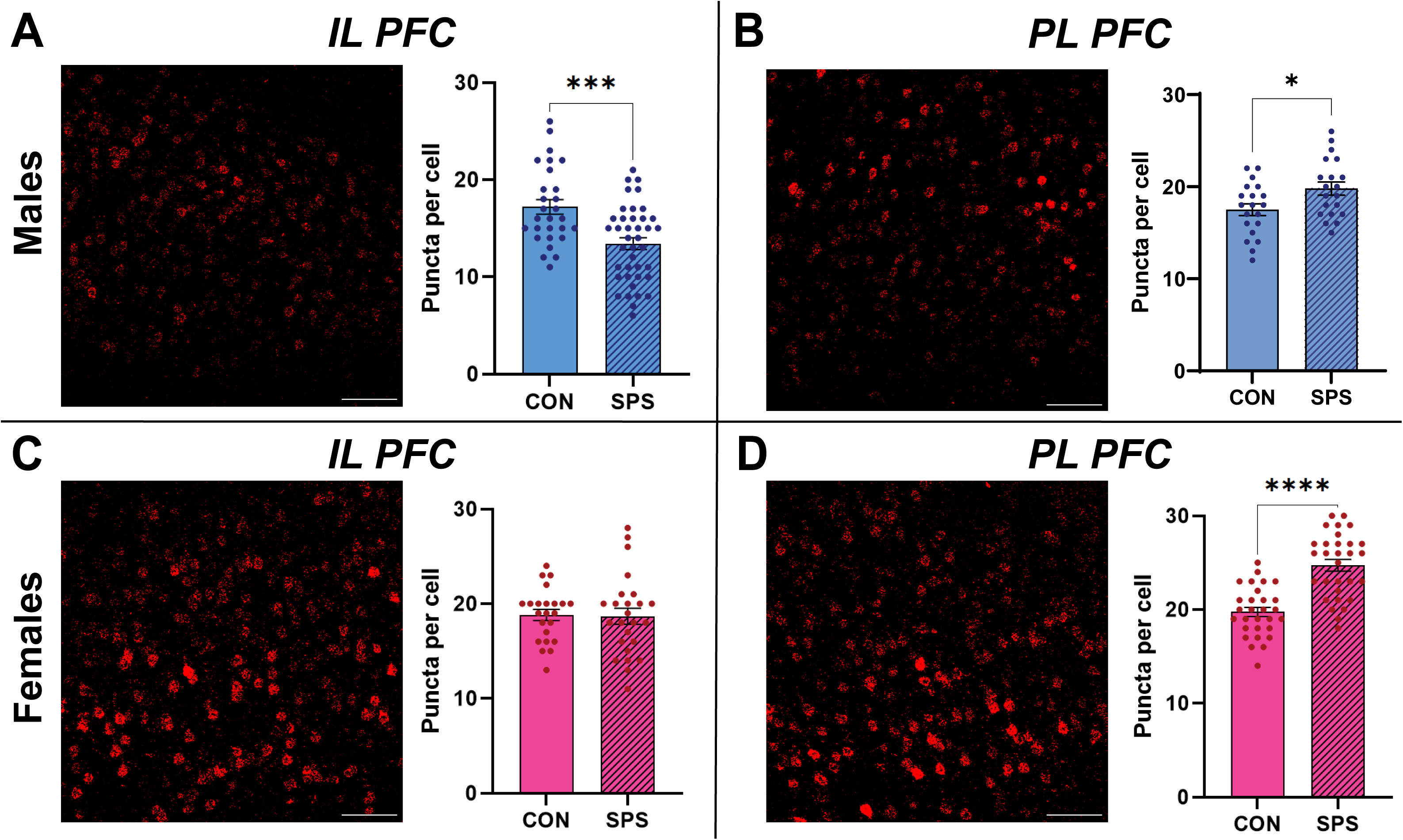
SPS Effects on PFC PACAP RNA Expression. (A) Male IL PACAP Expression. (B) Male PL PACAP Expression. (C) Female IL PACAP Expression. (D) Female PL PACAP Expression. N = 20-40 cells. Scale bar = 100 µm. * = p<0.05.

## 4. DISCUSSION

We utilized SPS and IL-targeted pharmacology to explore the role of PACAP and sex in PTSD-like phenotypes. These studies revealed differential effects within sexes following SPS, as well as a potential role for IL PACAP signaling in governing this divergence in stress resilience.

PTSD disproportionately impacts women, often yielding unique symptoms, making the study of sex differences key to understanding this disorder^18,31–34^. For example, most people experiencing PTSD will exhibit symptoms of re-experiencing and avoidance of cues related to the trauma, yet men tend to show more arousal-related symptoms and women tend to show more mood and cognition-related symptoms. Women also generally exhibit heightened symptom severity^18,23,31,35^. On the other hand, SPS in rats typically elicits stronger behavioral phenotypes in males, impacting domains associated with fear, anxiety, arousal, and sociability^3,22,36,37^. While females do show some dysregulation in fear conditioning reinstatement and avoidance, these behaviors are often more subtle^17,24,38^. This may be due to the fact that the SPS paradigm was developed in and validated with behavioral tests designed for males^3,22,36^. However, SPS does impact females on molecular level, often inducing changes in Fos and glucocorticoid receptor expression throughout the brain^17,23,24^, suggesting that it is still a valuable model for understanding how females react to severely stressful experiences.

Here, we were able to uncover female SPS phenotypes in the PTSD-relevant domains of anxiety (i.e., decreased OF center time), threat (i.e., increased threat avoidance and decreased threat assessment), and sociability (i.e., increased social interaction). Our ability to reveal these endpoints is primarily attributed to the approach of behavioral indexing, where instead of looking at individual behavioral endpoints, we examined broader behavioral domains by considering these individual endpoints in tandem. The heightened threat avoidance and blunted threat assessment phenotypes revealed with this method aligns nicely with PTSD symptomology in both men and women^2,22,35,36^. The negative correlation between threat avoidance and appraisal also suggests individual differences within SPS severity and coping strategies, another feature often seen in humans suffering from PTSD^2,3^. An additional key finding here is the variance by sex in the sociability domain, with males having decreased sociability where females had increased sociability. This would suggest that where males may be more likely to withdraw from conspecifics, females may be more likely to seek out social peers. In humans, women are more likely to engage socially after trauma as a coping mechanism than men^31,35,39^. In fact, women who do not seek social support tend to have worse PTSD outcomes^31,35^, a notion that is supported here by a negative correlation between threat avoidance and sociability in female SPS rats. These findings not only show the value of behavioral indexing, but also have interesting implications for behavioral PTSD therapies in men vs. women.

We chose to focus on IL PACAP due to the established roles of the human vmPFC (analog to rodent IL) in governing PTSD symptoms^12,17,33,40^, along with known sex differences in PFC PACAP^6–10^. Using intra-IL cannulae, a PACAP agonist or antagonist was infused directly into the IL prior to SPS. While the resulting phenotypes were not overtly strong, there are some patterns that are consistent with a role for IL PACAP in SPS, as well as differences by sex in PACAP signaling. Overall, it appears that IL ANTAG improved SPS phenotypes in males, while IL PACAP worsened SPS phenotypes in females. There is some important background information for interpreting these results in the present context. First, the IL-amygdala connection plays an important role in fear and anxiety-like behaviors^12,17,41^. Typically, the IL inhibits the CEA via BLA connections, blunting expression of these phenotypes (Fig 7A).

**Figure 7:**
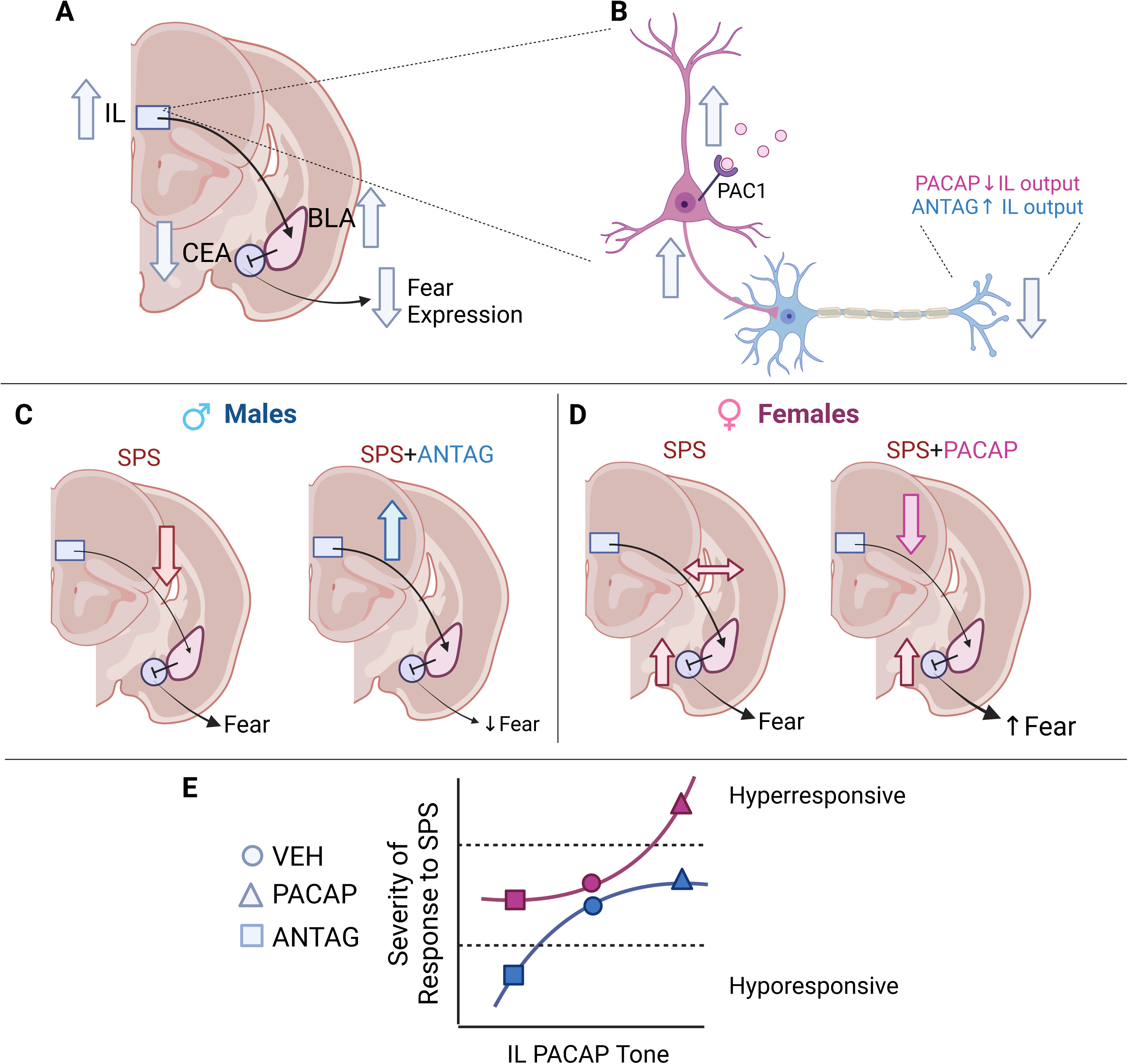
Summary of the Interactions between SPS and PACAP in Males and Females. (A) The IL sends excitatory projections to the BLA, which inhibits the CEA via interneuron connections. This decreased CEA output suppresses fear expression. Thus, increased IL activity would typically decrease fear via this IL-amygdala circuit. (B) Within the IL, PAC1 receptors are primarily on inhibitory interneurons, suggesting that an increase in PACAP in this region would increase inhibitory activity, ultimately decreasing IL output. ANTAG would have the opposite effect, increasing IL output. (C) Previously our lab has demonstrated that, in males, SPS decreases IL excitation of the BLA, yielding fear expression in this model. In this case, infusing ANTAG into the IL restores this IL-BLA connection (via decreased interneuron activation), partly blocking the SPS effects observed here. (D) However, in females, SPS does not impact the IL-BLA connection, likely acting more downstream at the CEA level. In this case, increasing IL PACAP weakens the IL-BLA connection (via enhanced interneuron activation), enhancing SPS effects. (E) Note that the lack of effects in males receiving PACAP and females receiving ANTAG suggests a healthy range of IL PACAP tone, along with differences by sex within SPS x PACAP interactions. It appears that the male response to SPS (blue) is not impacted by increasing PACAP but can be blunted in cases where PACAP tone is sufficiently decreased (ANTAG). On the other hand, the female response to SPS (pink) is not impacted by decreasing PACAP but can be exacerbated in cases where PACAP tone is sufficiently increased (PACAP). These unique male and female effects in pharmacology and circuitry have intriguing implications for not only how we understand the impact of neuropeptides on anxiety-like behaviors following traumatic stress, but also potential treatment options for PTSD in humans.

Second, the main PACAP receptors (PAC1 receptors) are primarily expressed on inhibitory interneurons in the IL^12^. Therefore, increasing PACAP in this region would increase inhibitory drive, ultimately decreasing the excitatory output from the IL (Fig 7B). Finally, this fear-related circuitry is differentially impacted by SPS in males and females^17^. In males, SPS decreases IL output, weakening the IL-BLA connection, permitting overall increased CEA output to drive SPS-related fear and anxiety-like phenotypes^17,19,42^ (Fig 7C). However, different circuitry alterations seem to drive SPS phenotypes in females^17^. We have previously proposed that the IL-BLA activity remains intact, while the CEA, this circuit’s ultimate output node, instead exhibits direct hyperactivation^17^ (Fig 7D). Thus, female SPS phenotypes emerge further downstream than male phenotypes. Considering these points together lends insight into the mechanism governing the present PACAP results. We propose that, in males, administering ANTAG to the IL would be expected to increase IL output via disinhibition of pyramidal neurons (Fig 7B). This output increase counteracts the SPS-induced decrease in IL output^42^, reducing overall amygdala output, helping to block SPS phenotypes (Fig 7C). On the other hand, administering PACAP to females instead mimics decreased IL-BLA connectivity usually seen in males subjected to SPS, via increased interneuron activity in the IL (Fig 7B). PACAP thus exacerbates SPS phenotypes in females by adding onto the preexisting hyperactivation of the CEA (Fig 7D). Thus, underlying sex differences in PACAP physiology and SPS circuitry help to explain the divergent actions of IL PACAP manipulations by sex in the present studies.

A remaining question with this interpretation is why PACAP had minimal effects in males and ANTAG had minimal effects in females. It is important to remember that PACAP is a neuropeptide that acts to fine-tune circuitry^4,5,11,43,44^. There is an inherent inverted-U to this type of molecule, with an optimal range of PACAP tone that helps to maintain normal function. - We posit that inherent sex differences within this U-shaped curve contribute to the present results (Fig 7E). The relationship between PACAP tone and SPS responsivity may be shifted in each sex, with females having higher sensitivity to increased PACAP (Fig 7E, pink line) and males having higher sensitivity to decreased PACAP (Fig 7E, blue line). Such a relationship would mean that females are less subject to stress-ameliorating effects of low PACAP but more vulnerable to stress-promoting effects of high PACAP. Therefore, in females, the PACAP agonist was sufficient to increase SPS severity (Fig 7E, pink triangle), but ANTAG was unable to lower the levels enough to offer much benefit (Fig 7E, pink square). On the other hand, males, with greater SPS responses to PACAP decreases than increases, were protected from harmful effects of too much PACAP (Fig 7E, blue triangle) and benefited from ANTAG’s effects (Fig 7E, blue square). Thus, most of the animals in the present study fall within the optimal range of IL PACAP, including females receiving ANTAG and males receiving PACAP, and exhibit typical SPS effects. It is only in the PACAP females and ANTAG males that we move far enough along these differential curves to elicit heighted SPS effects in females and blunted SPS effects in males.

The present RNAscope results offer further insight into these relationships, namely that the IL may not be the most important PFC region for females with regard to SPS. Males exhibit decreased IL PACAP mRNA one week after SPS. This decrease in males may be compensatory, with the IL downshifting its production of PACAP in response to an initial increase acutely following SPS. While future studies examining acute responses to SPS are warranted, this supports that idea that males may benefit from IL ANTAG, as this may counteract an initial increase in PACAP following SPS. On the other hand, females show no change in IL PACAP mRNA after SPS. This could mean the female IL does not inherently respond as much to SPS as the male IL, a result which aligns with our lab’s previous SPS FOS data^17^.

Instead, the females show robust changes in the adjacent PL, with PACAP mRNA increasing one week after SPS. Where the IL inhibits amygdala output, the PL typically drives it, promoting defensive and stress-like behaviors^41,45,46^. An increase in PL PACAP following SPS, therefore, would be expected to serve a protective effect, inhibiting the PL via enhanced interneuron activation, thereby inhibiting the amygdala and its stress-promoting effects. The PL has also been associated with PACAP sex differences, with females responding more to PL PACAP manipulations than males on fear conditioning endpoints^20^. While males show a PL PACAP mRNA increase in the present study as well, this change is to a smaller degree. Thus, the PL may be an intriguing future target for PACAP manipulations during SPS, especially in females.

Another interesting future direction for these studies would be examining SPS effects on other PTSD-related behavioral domains in males and females. We have examined fear conditioning in the past and found that males are more susceptible to SPS-related extinction deficits, while females have stronger contextual and reinstatement phenotypes^17^. PTSD is also associated with impairments in cognition, motivation, and arousal^2,47,48^. SPS is known, at least in males, to impact similar behaviors, including spatial learning in the Morris water maze, attentional set shifting, sucrose preference, and acoustic startle^25,36^. Given the divergent social effects observed here, it would be interesting to examine SPS (and PACAP) effects on these behaviors in females, as well as on female exclusive behaviors such as maternal care^49^.

Further exploring the molecular components of these phenotypes would also be valuable. One element of this would be studying these relationships in other stress-regulating regions^4,13,41^.

PACAP manipulation alone in the PL, BST, and BLA has been shown to impact fear- and anxiety-like behaviors^11,20,21,27^. It would be interesting to see if SPS effects in these regions are similarly affected by PACAP. Another element would be a deeper examination of cell and receptor types. PACAP acts on multiple receptor subtypes, and while the target of these studies, the PAC1 receptor, has the highest affinity, PACAP also acts on VPAC1 and VPAC2 receptors which contribute to its function^4,11,13,50^. These receptors are expressed on various cell types. While this is mostly interneurons in the PFC^12^, PACAP receptors are present in low levels on pyramidal neurons and glia which may also play a role in the present SPS phenotypes^11,51–54^. Conducting studies on which elements in the PACAP system contribute to which SPS phenotypes is an important next step in understanding the complex relationships explored here.

## 5. CONCLUSIONS

These studies lend valuable insight into the interplay between SPS, sex, and PACAP which could have important implications for our understanding of PTSD. The unique male and female SPS behavioral phenotypes noted here are similar to PTSD symptomologies observed in men and women, namely congruent threat-related behaviors accompanied by divergent social characteristics. Male and female fear circuits exhibit differential responses to SPS, as well as IL PACAP, resulting in divergent effects of PACAP agonism and antagonism, with males showing improvement with ANTAG and females showing impairment with PACAP. Overall, the present results suggest that SPS can inform how PTSD impacts different behavioral domains in males and females, and that PACAP is an intriguing candidate contributing to these differences.

## Author contributions

SEM, RDM, EMC, and JPH conceptualized the study together. JBC, BAP, SEM, RDM, EMC, and MAS ran the behavioral experiments. CM and BAP assessed cannula placement. BAP ran the HPA axis analyses. RKP ran the RNAscope analyses. JBC and MAS analyzed the behavioral and molecular data. MAS, JPH, NN, and RKP participated in interpreting the results. MAS generated figures and wrote the manuscript. MAS and JPH edited the manuscript. All authors participated in finalizing the manuscript.

## Acknowledgments

The authors would like to thank Rachel Morano, Parinaz Mahbod, and the Herman lab staff for their assistance with animal experiments.

## Data Availability

All data are available in the main text or supplementary materials.

## ABBREVIATIONS

BLA: Basolateral Amygdala
BST: Bed Nucleus of the Stria Terminalis
CEA: Central Amygdala
CORT: Corticosterone
EPM: Elevated Plus Maze
HPA: Hypothalamic-Pituitary-Adrenal
IL: Infralimbic Prefrontal Cortex
NOR: Novel Object Recognition
OF: Open Field
PAC1: PACAP Receptor
PACAP: Pituitary Adenylate Cyclase-Activating Polypeptide
PFC: Prefrontal Cortex
PL: Prelimbic Prefrontal Cortex
PTSD: Post-Traumatic Stress Disorder
SI: Social Interaction
SPS: Single Prolonged Stress
vmPFC: Ventromedial Prefrontal Cortex

## BIBLIOGRAPHY

1. Schein, J. et al. Prevalence of post-traumatic stress disorder in the United States: a systematic literature review. Current Medical Research and Opinion 37, 2151–2161 (2021).

2. 2. NIH. Post-Traumatic Stress Disorder. https://www.nimh.nih.gov/health/topics/post-traumatic-stress-disorder-ptsd (2023).

3. Lisieski, M. J., Eagle, A. L., Conti, A. C., Liberzon, I. & Perrine, S. A. Single-prolonged stress: A review of two decades of progress in a rodent model of post-traumatic stress disorder. Frontiers in Psychiatry 9, 1–22 (2018).

4. Hammack, S. E. & May, V. Pituitary adenylate cyclase activating polypeptide in stress-related disorders: Data convergence from animal and human studies. Biological Psychiatry 78, 167–177 (2015).

5. Mustafa, T. Pituitary adenylate cyclase-activating polypeptide (PACAP): a master regulator in central and peripheral stress responses. *Advances in pharmacology (San Diego*, Calif.) 68, 445–57 (2013).

6. Almli, L. M. et al. ADCYAP1R1 genotype associates with post-traumatic stress symptoms in highly traumatized African-American females. *American Journal of Medical Genetics*, Part B: Neuropsychiatric Genetics 162, 262–272 (2013).

7. Jovanovic, T. et al. PAC1 receptor (ADCYAP1R1) genotype is associated with dark- enhanced startle in children. Molecular Psychiatry 18, 742–743 (2013).

8. Ressler, K. J. et al. Post-traumatic stress disorder is associated with PACAP and the PAC1 receptor. Nature 470, 492–497 (2011).

9. Liao, C. et al. Targeting the PAC1 Receptor for Neurological and Metabolic Disorders. Current Topics in Medicinal Chemistry 19, 1399–1417 (2019).

10. Uddin, M. et al. ADCYAP1R1 genotype, posttraumatic stress disorder, and depression among women exposed to childhood maltreatment. Depression and Anxiety 30, 251–258 (2013).

11. King, S. B., Toufexis, D. J. & Hammack, S. E. Pituitary adenylate cyclase activating polypeptide (PACAP), stress, and sex hormones. Stress 20, 465–475 (2017).

12. Martelle, S. E. et al. Prefrontal cortex PACAP signaling: organization and role in stress regulation. Stress 24, 196–205 (2021).

13. Boucher, M. N., May, V., Braas, K. M. & Hammack, S. E. PACAP orchestration of stress-related responses in neural circuits. Peptides 142, 170554 (2021).

14. Lehmann, M. L., Mustafa, T., Eiden, A. M., Herkenham, M. & Eiden, L. E. PACAP-deficient mice show attenuated corticosterone secretion and fail to develop depressive behavior during chronic social defeat stress. Psychoneuroendocrinology 38, 702–715 (2013).

15. Morita, Y. et al. Lack of trimethyltin (TMT)-induced elevation of plasma corticosterone in PACAP-deficient mice. Annals of the New York Academy of Sciences 1070, 450–456 (2006).

16. Hattori, S. et al. Comprehensive behavioral analysis of pituitary adenylate cyclase-activating polypeptide (PACAP) knockout mice. Frontiers in Behavioral Neuroscience 6, 1–18 (2012).

17. Cotella, E. M. et al. Adolescent Stress Confers Resilience to Traumatic Stress Later in Life: Role of the Prefrontal Cortex. Biological Psychiatry Global Open Science 3, 274–282 (2023).

18. Ramikie, T. S. & Ressler, K. J. Mechanisms of Sex Differences in Fear and Posttraumatic Stress Disorder. Biological Psychiatry 83, 876–885 (2018).

19. Marek, R., Sun, Y. & Sah, P. Neural circuits for a top-down control of fear and extinction. Psychopharmacology 236, 313–320 (2019).

20. Kirry, A. J. et al. Pituitary adenylate cyclase-activating polypeptide (PACAP) signaling in the prefrontal cortex modulates cued fear learning, but not spatial working memory, in female rats. Neuropharmacology 133, 145–154 (2018).

21. Cho, J. H. et al. Pituitary adenylate cyclase-activating polypeptide induces postsynaptically expressed potentiation in the intra-amygdala circuit. Journal of Neuroscience 32, 14165– 14177 (2012).

22. Souza, R. R., Noble, L. J. & McIntyre, C. K. Using the Single Prolonged Stress Model to Examine the Pathophysiology of PTSD. Frontiers in pharmacology 8, 615 (2017).

23. Pooley, A. E. et al. Sex differences in the traumatic stress response: PTSD symptoms in women recapitulated in female rats. Biology of Sex Differences 9, (2018).

24. Keller, S. M., Schreiber, W. B., Staib, J. M. & Knox, D. Sex differences in the single prolonged stress model. Behavioural Brain Research 286, 29–32 (2015).

25. Mancini, G. F. et al. Sex-divergent long-term effects of single prolonged stress in adult rats. Behavioural Brain Research 401, 113096 (2021).

26. Cotella, E. M. et al. Lasting Impact of Chronic Adolescent Stress and Glucocorticoid Receptor Selective Modulation in Male and Female Rats. Psychoneuroendocrinology 112, 104490 (2020).

27. Lezak, K. R. et al. Pituitary adenylate cyclase-activating polypeptide (PACAP) in the bed nucleus of the stria terminalis (BNST) increases corticosterone in male and female rats. Psychoneuroendocrinology 45, 11–20 (2014).

28. Telegdy, G. & Kokavszky, K. The action of pituitary adenylate cyclase activating polypeptide (PACAP) on passive avoidance learning. the role of transmitters. Brain Research 874, 194– 199 (2000).

29. Paxinos, G. & Charles Watson. The Rat Brain in Stereotaxic Coordinates Sixth Edition. Elsevier Academic Press (2007).

30. Maren, S. & Holmes, A. Stress and Fear Extinction. Neuropsychopharmacol **41**, 58–79 (2016).

31. Eder-Moreau, E. et al. Neurobiological Alterations in Females With PTSD: A Systematic Review. Frontiers in Psychiatry 13, (2022).

32. Shansky, R. M. Sex differences in PTSD resilience and susceptibility: Challenges for animal models of fear learning. Neurobiology of Stress 1, 60–65 (2015).

33. Bangasser, D. A. & Valentino, R. J. Sex differences in stress-related psychiatric disorders: Neurobiological perspectives. Frontiers in Neuroendocrinology 35, 303–319 (2014).

34. 34. PTSD: National Center for PTSD. *U.S. Department of Veterans Affairs* https://www.ptsd.va.gov/understand/common/common_adults.asp (2018).

35. Hu, J. et al. Gender Differences in PTSD: Susceptibility and Resilience. Gender Differences in Different Contexts (2017) doi:10.5772/65287.

36. Yamamoto, S. et al. Single prolonged stress: Toward an animal model of posttraumatic stress disorder. Depression and Anxiety 26, 1110–1117 (2009).

37. Eagle, A. L., Fitzpatrick, C. J. & Perrine, S. A. Single prolonged stress impairs social and object novelty recognition in rats. Behavioural Brain Research 256, 591–597 (2013).

38. Collins, B. et al. The role of avoidance in modulating single prolonged stress effects on emotional memory in male and female rats. Behavioural brain research 452, 114579 (2023).

39. Olff, M. Sex and gender differences in post-traumatic stress disorder: an update. European Journal of Psychotraumatology 8, 4 (2017).

40. Stark, E. A. et al. Post-traumatic stress influences the brain even in the absence of symptoms: A systematic, quantitative meta-analysis of neuroimaging studies. Neuroscience and Biobehavioral Reviews 56, 207–221 (2015).

41. Ulrich-Lai, Y. M. & Herman, J. P. Neural regulation of endocrine and autonomic stress responses. Nature Reviews Neuroscience 10, 397–409 (2009).

42. Nawreen, N., Baccei, M. L. & Herman, J. P. Single Prolonged Stress Reduces Intrinsic Excitability and Excitatory Synaptic Drive Onto Pyramidal Neurons in the Infralimbic Prefrontal Cortex of Adult Male Rats. Front. Cell. Neurosci. 15, (2021).

43. Bucher, E. A. et al. Coherence and cognition in the cortex: the fundamental role of parvalbumin, myelin, and the perineuronal net. Brain Structure and Function 226, 2041– 2055 (2021).

44. Gilmartin, M. R. & Ferrara, N. C. Pituitary Adenylate Cyclase-Activating Polypeptide in Learning and Memory. Frontiers in Cellular Neuroscience 15, 663418 (2021).

45. Knox, D. et al. Neural circuits via which Single prolonged stress exposure leads to fear extinction retention deficits. Learning and Memory 23, 689–698 (2016).

46. Giustino, T. F. & Maren, S. The role of the medial prefrontal cortex in the conditioning and extinction of fear. Frontiers in Behavioral Neuroscience 9, (2015).

47. Bangasser, D. A., Eck, S. R. & Ordoñes Sanchez, E. Sex differences in stress reactivity in arousal and attention systems. Neuropsychopharmacology 44, 129–139 (2019).

48. Donahue, R. J., Venkataraman, A., Carroll, F. I., Meloni, E. G. & Carlezon, W. A. Pituitary Adenylate Cyclase–Activating Polypeptide Disrupts Motivation, Social Interaction, and Attention in Male Sprague Dawley Rats. Biological Psychiatry 80, 955–964 (2016).

49. Ivy, A. S., Brunson, K. L., Sandman, C. & Baram, T. Z. Dysfunctional nurturing behavior in rat dams with limited access to nesting material: a clinically relevant model for early-life stress. Neuroscience 154, 1132–1142 (2008).

50. Hirabayashi, T., Nakamachi, T. & Shioda, S. Discovery of PACAP and its receptors in the brain. The journal of headache and pain 19, (2018).

51. Figiel, M. & Engele, J. Pituitary Adenylate Cyclase-Activating Polypeptide (PACAP), a Neuron-Derived Peptide Regulating Glial Glutamate Transport and Metabolism. The Journal of Neuroscience 20, 3596 (2000).

52. Kong, L. et al. Pituitary Adenylate cyclase-activating polypeptide orchestrates neuronal regulation of the astrocytic glutamate-releasing mechanism system xc (.). Journal of neurochemistry 137, 384–393 (2016).

53. Taylor, R. D. T. et al. Pituitary adenylate cyclase-activating polypeptide (PACAP) inhibits the slow afterhyperpolarizing current sIAHP in CA1 pyramidal neurons by activating multiple signaling pathways. Hippocampus 24, 32 (2014).

54. Zhang, L. et al. Behavioral role of pacap signaling reflects its selective distribution in glutamatergic and gabaergic neuronal subpopulations. eLife 10, 1–77 (2021).

