## Supplementary Information and Supplementary Figures for "Sex-Specific Regulation of Behavioral Responses to Single Prolonged Stress: Role of PACAP"

Marissa A. Smail^1,2,3, #^, Evelin M. Cotella^1,4^, Susan E. Martelle^1,5^, James B. Chambers^1^, Ria K. Parikh^1,6^, Christine E. Moore^1^, Ben A. Packard^1^, Nawshaba Nawreen^1,7^, Rachel D. Moloney^1,8,9,10^, James P. Herman^1,11,12^

**Table of Contents**

**Supplementary Figures**

- **Supplementary Figure 1**: Additional Behavioral Results
- **Supplementary Figure 2**: Corticosterone Response Curves
- **Supplementary Figure 3**: Behavior Index Correlations

**Supplementary Table** (as *separate Excel file*):

- **Supplementary Table 1**: Detailed Statistical Results

**SUPPLEMENTARY FIGURES**

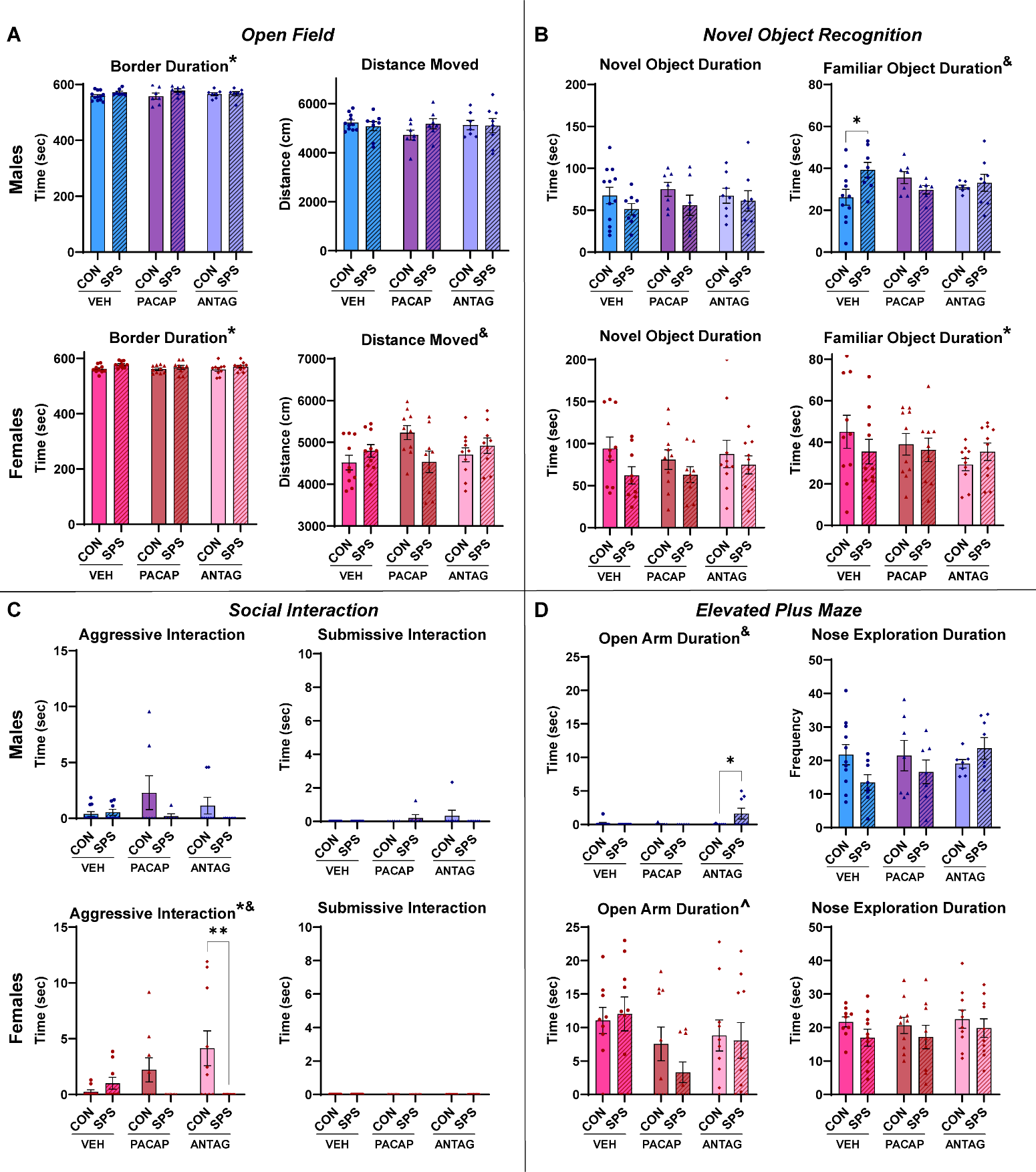

**Supplementary Figure 1: Additional Behavioral Results.** (A) Open Field border duration and distance moved. (B) Novel Object novel and familiar object duration. (C) Social Interaction aggressive and submissive interaction duration. (D) Elevated Plus Maze open arm duration and nose exploration duration. n = 7-12. Blue = males. Pink = females.
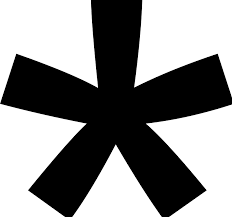
 = Main Effect of Stress. **^** = Main Effect of Infusion. **&** = Interaction. * = post hoc p<0.05.

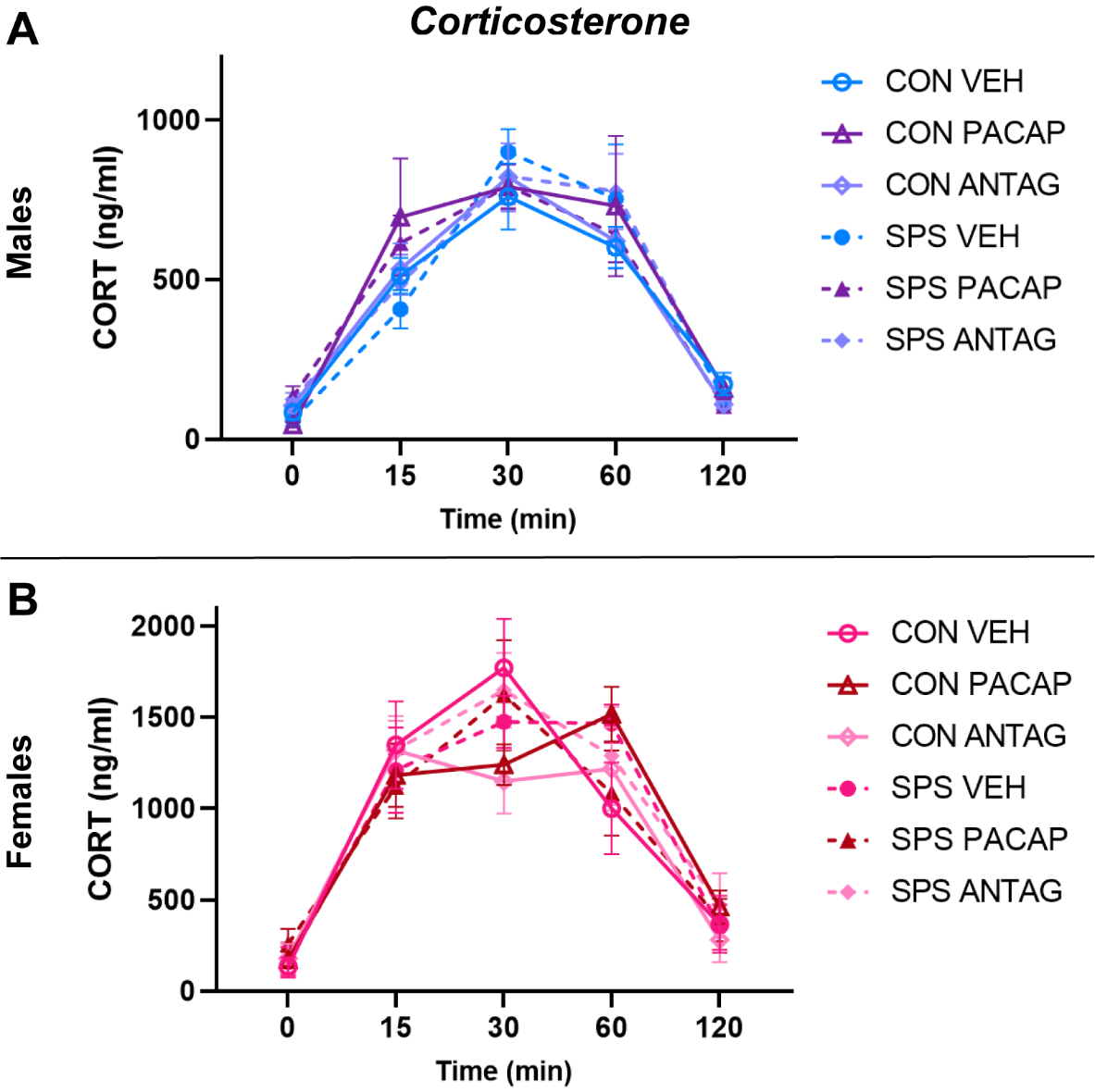

**Supplementary Figure 2: Corticosterone Response Curves.** (A) Male CORT response to 30 min restraint. (B) Female CORT response to 30 min restraint. n = 5-12.

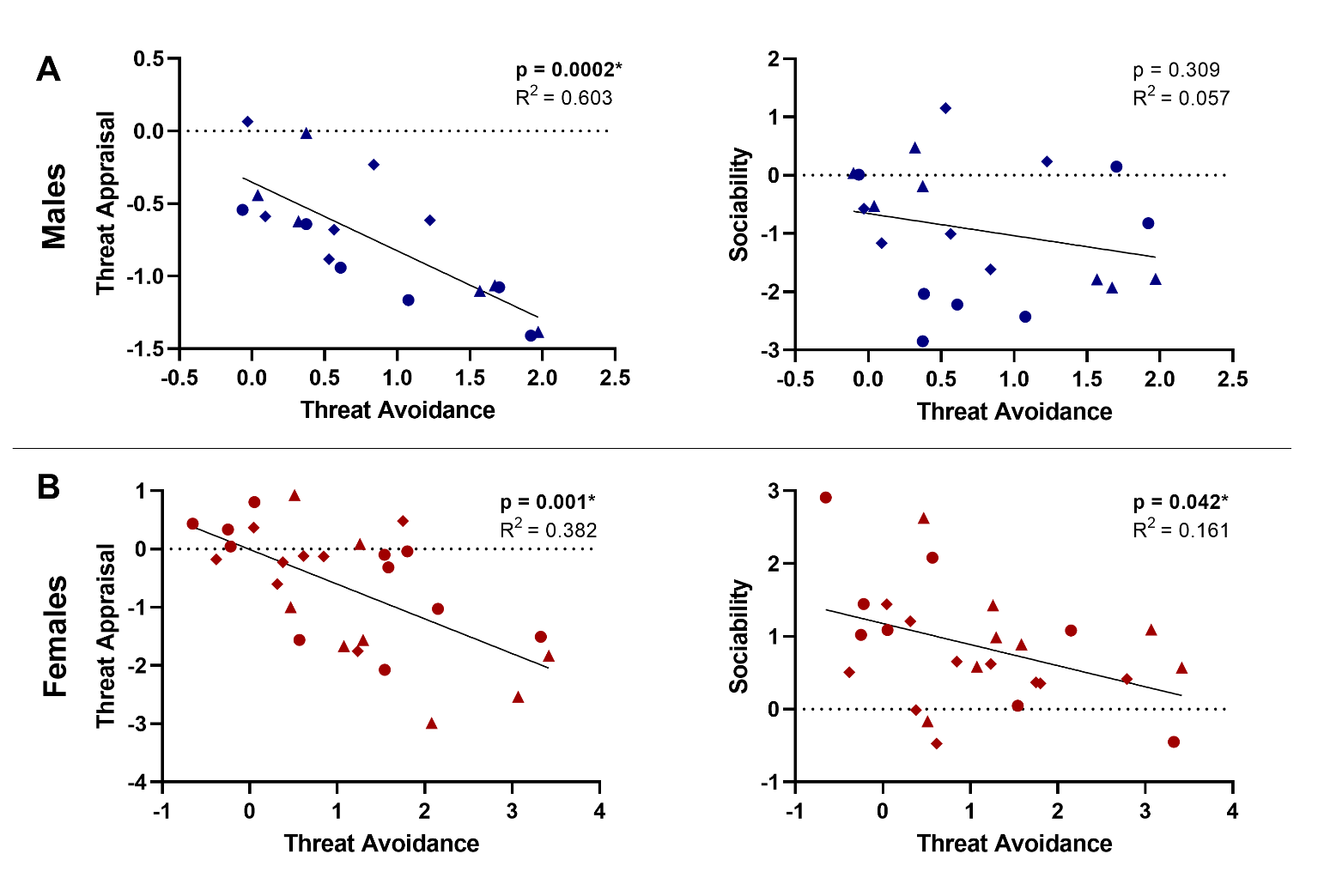

**Supplementary Figure 3: Behavior Index Correlations.** (A) Correlations between threat avoidance, threat appraisal, and sociability in SPS males. (B) Correlations between threat avoidance, threat appraisal, and sociability in SPS females. n = 20-26. Circle = VEH. Triangle = PACAP. Diamond = ANTAG.

**SUPPLEMENTARY TABLE (as separate Excel file)**

**Supplementary Table 8: Detailed Statistical Results.**

*Tab 1: Main Body Figures*. Statistical results for Figures 2-6.

*Tab 2: Supplementary Figures*. Statistical results for Supplementary Figures 1-2.
